# The Regulatory Mendelian Mutation score for GRCh38

**DOI:** 10.1101/2022.03.14.484240

**Authors:** Lusiné Nazaretyan, Martin Kircher, Max Schubach

**Affiliations:** Berlin Institute of Health at Charité – Universitätsmedizin Berlin, Berlin, Germany; Institute of Human Genetics, University of Lübeck, Lübeck, Germany

## Abstract

**Motivation:** Various genome sequencing efforts for individuals with rare Mendelian disease have increased the research focus on the non-coding genome and the clinical need for methods that prioritize potentially disease causal non-coding variants. Some methods and annotations are not available for the current human genome build (GRCh38), for which the adoption in databases, software and pipelines was slow.

**Results:** Here, we present an updated version of the Regulatory Mendelian Mutation (ReMM) score, re-trained on features and variants derived from the GRCh38 genome build. Like its GRCh37 version, it achieves good performance on its highly imbalanced data. To improve accessibility and provide users with a toolbox to score their variant files and lookup scores in the genome, we developed a website and API for easy score lookup.

**Availability and Implementation:** Pre-scored whole genome files of GRCh37 and GRCh38 genome builds are available on Zenodo https://doi.org/10.5281/zenodo.6576087. The website and API are available at https://remm.bihealth.org.

## INTRODUCTION

The Regulatory Mendelian Mutation (ReMM) score predicts the potential pathogenicity of non-coding variations in the human reference genome build GRCh37/hg19 (Smedley et al. 2016). The updated reference genome GRCh38/hg38 contains new sequences at nearly 100 assembly gaps and reduces unresolved bases at about 3% of the genome (Guo et al. 2017). This establishes a need for an update of the ReMM score and we present a version developed particularly for GRCh38. Further, we update the ReMM score for GRCh37 by including feature updates and an improvement in handling missing values. Finally, we provide a webserver and API for scoring VCF files, single variant lookups or range lookups.

## IMPLEMENTATION

The ReMM score is based on an imbalanced-aware machine learning algorithm, hyperSMURF (Schubach et al. 2017), trained from known pathogenic non-coding variants of Mendelian disorders and a set of putatively benign variants. As pathogenic set, we use 406 hand-curated variants already used in the prior version (Smedley et al. 2016), lifted to GRCh38 using UCSC liftOver (v377) (Lee et al. 2022). The proxy-benign set includes around 14 million of human-lineage-derived sequence alterations (Rentzsch et al. 2019), which we filtered to non-coding sequence using Jannovar v0.36 (Jäger et al. 2014) and RefSeq (O’Leary et al. 2016). The high imbalance after non-coding filtering is similar on both genome builds (14.8M and 13.9M negatives for GRCh37 and GRCh38, respectively). Therefore, we kept parameters for hyperSMURF as determined in Smedley et al. 2016 (Supplementary Table 1). Because of the large and computationally expensive dataset, we replaced hyperSMURF with the updated parSMURF implementation (Petrini et al. 2020).

Genomic data is confounded by local correlation of annotations. Further, known pathogenic variants are not distributed evenly across the genome (e.g., due to selection bias). When not accounted for, learners might infer superior hold-out performance because of genomic proximity of variants. To handle the local correlation structure, we apply cytogenic band-aware cross-validation using ten folds (Smedley et al. 2016).

Twenty-six selected features (see Supplementary Table 2) capture functional constraint and sequence functions (sequence composition, epigenetics, conservation, population variance and regulatory regions). The feature set was kept close to the original feature set of ReMM, but some were not available from the original databases or were updated. Some features have a high proportion of missing values and the initial version of ReMM imputed all of them with zero. In genomics, a missing value often indicates an experimental signal that is too low to be measured, in line with this imputation. We have now identified some features (e.g., GC content or conservation scores) where the genome-wide average of the annotation is more appropriate and impute them differently in this version (see Supplementary Table 2). For missing p-values, we use the value 1.

Pre-scored, block-gzip compressed and indexed whole genome files (Li 2011) were generated to allow a fast scoring of variants as well as an easy integration into other software. Every genomic position was scored with a general ReMM model trained on all data (v0.4.hg19 and v0.4.hg38, respectively). To guarantee unbiased score usage, e.g., for performance benchmarks with other tools, we replaced the score of variants in the training set with the cross-validated scores. The training and scoring pipeline is implemented in snakemake, a workflow management system for reproducible and scalable analysis (Mölder et al. 2021).

## RESULTS

### Performance of ReMM on GRCh38

After 100 training cycles using different random seeds and ten-fold cytoband cross validation, we achieve an excellent performance with an average area under the precision recall curve (AUPRC) of 0.613±0.005 (Supplementary Table 3). We randomly picked one model for the final scoring with an AUPRC of 0.610 (Supplementary Figure 1a, ROC performance available in Supplementary Table 1b).

Rather than using ReMM scores for ranking, some users need to specify score thresholds for classifying into pathogenic and benign variants. Using a cutoff of 0.5 yields a good result in terms of retrieving known pathogenic non-coding variants (i.e., recall or True Positive rate, TP), but the number of negatives might be extremely large. For ReMM v0.4.hg38, recall is 92% (375 out of 406) at a cutoff of 0.5 (Supplementary Figure 1c), but precision is close to zero with lots of false positives (FP) (86,507 out of 13,911,061; FP rate=0.006). The F1-score (harmonic mean of recall and precision) is highest at 0.963, resulting in a TP rate of 0.554 and a FP rate of 5.3e-6. Using the F2-score, we can give more weight to recall. Here, the optimal cutoff is 0.914, resulting in a TP rate of 0.702 and a FP rate of 2.3e-5. Analogous to NCBI ClinVar (Landrum et al. 2018) pathogenic and likely pathogenic categories, we suggest to use a ReMM score above the F1 threshold as weak computational evidence for “pathogenic” and a score above the F2 threshold and below the F1 threshold for “likely pathogenic”. For ReMM v0.4.hg19, these thresholds are 0.961 and 0.924 (Supplementary Figure 1d), respectively.

### Correlation of scores and features

To compare both genome builds, we correlate ReMM scores from three genomic regions (genic content and not overlapping with assembly gap changes) and 120K randomly sampled positions and find that scores are highly correlated between versions (Supplementary Table 4). We also used these regions and sites to explore the average feature correlation (Supplementary Table 5). Further, we compare feature correlations between the genome builds directly on the training data (Supplementary Figure 2). As expected, we have highest correlation for sequence features, like GC content. Further, population variance features correlate well, with reduced correlation for the rare variant feature. This is likely due to spurious calls highly depending on the caller and the quality of the reference genome. We only see low correlation on sparse data (Fantom5).

### Imputing missing values

In previous ReMM versions, we used zero for missing values globally and trusted in the non-linearity of decision trees. Now, we use the average value of all defined positions for sequence and conservation features and one for p-values (see Supplementary Table 2). With the new approach, we see that the average AUPRC increases slightly (0.005 for v0.4.hg19, 0.009 for v0.4.hg38, Supplementary Table 6).

### Feature importance

From the underlying Ranger random forest (RF) models (Wright and Ziegler 2017), we can retrieve feature importance using the Gini index. We averaged values over all 100 RFs in the model (Supplementary Table 7). In general, mean feature importance scores distribute over all 26 features. No single feature stands out and our broad feature categories are all represented with at least one highly ranked feature. We interpret this as evidence that features were carefully picked and biases avoided. Epigenetic features increased in importance for the GRCh38 model (average rank 16 vs 19), maybe due to better mapping and processing of the underlying data. Fantom5 features are probably too sparse to receive high importance, but might be relevant for some variants. Between genome builds, feature importance values are similar and no significant change is detected (p-value 0.565, two-sided ranksum test). The replaced encRegTfbsClustered feature achieves a similar average Gini index (rank 6 on v0.4.hg19) as the previous numTFBSConserved feature (rank 4, data not shown).

## DATA AVAILABILITY

We precomputed the ReMM score for all sequence-resolved positions in the genome (GRCh37 and GRCh38 builds) and provide them on Zenodo (https://doi.org/10.5281/zenodo.6576087) or on our website (https://remm.bihealth.org). Our website also enables fast and easy scoring of variants. Variants can be uploaded via a VCF file (Danecek et al. 2011), or scores directly displayed with a single site or genomic range variant lookup. In addition, we provide a REST-API that allows tools and scripts to retrieve ReMM scores directly. Scoring on the website is available for both genome builds and all major ReMM versions.

## CONCLUSION

The ReMM v0.4 score is a fully retrained non-coding score available for both the GRCh37 and GRCh38 genome builds. Scores over the genome are highly correlated to the prior release with a performance increase due to the better coverage of features. In addition, we now established a reproducible and scalable framework for integration of new features or new training data for further development of ReMM. The pre-scored whole genome files and a website provide fast access and easy usage of the ReMM score for researchers in all areas. With this release, tools like Genomiser (Smedley et al. 2016) can now be run on the latest genome build, a highly demanded feature from the community.

