## Supplementary Figures and Tables for "The Regulatory Mendelian Mutation score for GRCh38"

### Supplementary Tables

**Supplementary Table 1: Hyperparameters** – Hyperparameters of ReMM training (all versions).

| Parameter | Value | Description |
|---|---|---|
| $n$ | 100 | Number of partitions |
| $o$ | 2 | SMOTE oversampling factor |
| $k$ | 5 | SMOTE k-nearest neighbor |
| $k$ | 3 | Undersampling factor |
| $t$ | 10 | Forest size for each Random Forest |
| $d$ | 5 | Random tree features for each Random Forest |

**Supplementary Table 2: Features** – Features used for training of the ReMM score v0.4 and their default missing value. The description column contains the original source from where features were downloaded.

| Feature | Missing value (hg19/hg38) | Description |
|---|---|---|
| CpGperGC | 65.085/ 65.8818 | Percentage of CpG island that is C or G.<br>https://hgdownload.soe.ucsc.edu/goldenPath/hg38/database/cpgIslandExt.txt.gz (hg19)<br>https://hgdownload.soe.ucsc.edu/goldenPath/hg38/database/cpgIslandExt.txt.gz (hg38) |
| CpGperCpG | 18.2498/ 18.7278 | Percentage of CpG island that is CpG.<br>https://hgdownload.soe.ucsc.edu/goldenPath/hg38/database/cpgIslandExt.txt.gz (hg19)<br>https://hgdownload.soe.ucsc.edu/goldenPath/hg38/database/cpgIslandExt.txt.gz (hg38) |
| CpGobsExp | 0.874298/ 0.874696 | Ratio of observed to expected CpGs in CpG island.<br>https://hgdownload.soe.ucsc.edu/goldenPath/hg38/database/cpgIslandExt.txt.gz (hg19)<br>https://hgdownload.soe.ucsc.edu/goldenPath/hg38/database/cpgIslandExt.txt.gz (hg38) |
| GCContent | 0.409034/ 0.408699 | GC-content in a window of ±75 bp of the reference genome |
| DnaseClusteredHyp | 0.0/0.0 | DnaseClustered (V3) hypersensitivity score<br>https://hgdownload.soe.ucsc.edu/goldenPath/hg19/database/wgEncodeRegDnaseClusteredV3.txt.gz (hg19)<br>https://hgdownload.soe.ucsc.edu/goldenPath/hg38/database/wgEncodeRegDnaseClustered.txt.gz (hg38) |
| DnaseClusteredScore | 0.0/0.0 | Number of DnaseClustered (V3) hypersensitive cells<br>https://hgdownload.soe.ucsc.edu/goldenPath/hg19/database/wgEncodeRegDnaseClusteredV3.txt.gz (hg19)<br>https://hgdownload.soe.ucsc.edu/goldenPath/hg38/database/wgEncodeRegDnaseClustered.txt.gz (hg38) |
| EncH3K27Ac | 0.0/0.0 | Maximum ENCODE H3K27 acetylation level<br>https://hgdownload.soe.ucsc.edu/gbdb/hg19/bbi/wgEncodeBroadHistone{Gm12878,H1hesc,Hsmm,Huvec,K562,Nhek,Nhlf}H3k27acStdSig.bigWig (hg19)<br>https://hgdownload.soe.ucsc.edu/gbdb/hg38/bbi/wgEncodeReg/wgEncodeRegMarkH3k27ac/wgEncodeBroadHistone{Gm12878,H1hesc,Hsmm,Huvec,K562,Nhek,Nhlf}H3k27acStdSig.bigWig (hg38) |
| EncH3K4Me1 | 0.0/0.0 | Maximum ENCODE H3K4 methylation level<br>https://hgdownload.soe.ucsc.edu/gbdb/hg19/bbi/wgEncodeBroadHistone{Gm12878,H1hesc,Hsmm,Huvec,K562,Nhek,Nhlf}H3k4me1StdSig.bigWig (hg19)<br>https://hgdownload.soe.ucsc.edu/gbdb/hg38/bbi/wgEncodeReg/wgEncodeRegMarkH3k27ac/wgEncodeBroadHistone{Gm12878,H1hesc,Hsmm,Huvec,K562,Nhek,Nhlf}H3k4me1StdSig.bigWig (hg38) |
| EncH3K4Me3 | 0.0/0.0 | Maximum ENCODE H3K4 methylation level<br>https://hgdownload.soe.ucsc.edu/gbdb/hg19/bbi/wgEncodeBroadHistone{Gm12878,H1hesc,Hsmm,Huvec,K562,Nhek,Nhlf}H3k4me3StdSig.bigWig (hg19)<br>https://hgdownload.soe.ucsc.edu/gbdb/hg38/bbi/wgEncodeReg/wgEncodeRegMarkH3k27ac/wgEncodeBroadHistone{Gm12878,H1hesc,Hsmm,Huvec,K562,Nhek,Nhlf}H3k4me3StdSig.bigWig (hg38) |
| Fantom5Perm | 0.0/0.0 | FANTOM 5 permissive enhancers<br>http://enhancer.binf.ku.dk/presets/permissive_enhancers.bed (hg19)<br>https://doi.org/10.5281/zenodo.545682 (hg38) |
| Fantom5Robust | 0.0/0.0 | FANTOM 5 robust enhancers<br>http://enhancer.binf.ku.dk/presets/robust_enhancers.bed (hg19)<br>https://doi.org/10.5281/zenodo.545682 (hg38) |
| encRegTfbsClustered | 0/0 | Number of ENCODE Regulation 'TF Clusters'<br>https://hgdownload.soe.ucsc.edu/goldenPath/hg19/database/encRegTfbsClustered.txt.gz (hg19) |

| | | |
|---|---|---|
| | | https://hgdownload.soe.ucsc.edu/goldenPath/hg38/database/encRegTfbsClustered.txt.gz (hg38) |
| priPhyloP | 0.0451375/ 0.0977878 | Primate PhyloP score<br>https://hgdownload.soe.ucsc.edu/goldenPath/hg19/phyloP46way/primates/ (hg19)<br>https://hgdownload.soe.ucsc.edu/goldenPath/hg38/phyloP17way/hg38.phyloP17way.wigFix.gz (hg38) |
| priPhastCons | 0.0977878/ 0.151415 | Primate PhastCons conservation score<br>https://hgdownload.soe.ucsc.edu/goldenPath/hg19/phastCons46way/primates/ (hg19)<br>https://hgdownload.soe.ucsc.edu/goldenPath/hg38/phastCons17way/hg38.phastCons17way.wigFix.gz (hg38) |
| verPhyloP | 0.0892683/ 0.0954381 | Vertebrate PhyloP score<br>https://hgdownload.soe.ucsc.edu/goldenPath/hg19/phyloP46way/vertebrate/ (hg19)<br>https://hgdownload.soe.ucsc.edu/goldenPath/hg38/phyloP100way/hg38.100way.phyloP100way/ (hg38) |
| verPhastCons | 0.102729/ 0.0980657 | Vertebrate PhastCons conservation score<br>https://hgdownload.soe.ucsc.edu/goldenPath/hg19/phastCons46way/vertebrate/ (hg19)<br>https://hgdownload.soe.ucsc.edu/goldenPath/hg38/phastCons100way/hg38.100way.phastCons/ (hg38) |
| mamPhyloP | 0.0357644/ 0.100913 | Mammalian PhyloP score.<br>https://hgdownload.soe.ucsc.edu/goldenPath/hg19/phyloP46way/placentalMammals/ (hg19)<br>https://hgdownload.soe.ucsc.edu/goldenPath/hg38/phyloP30way/hg38.30way.phyloP/ (hg38) |
| mamPhastCons | 0.0878941/ 0.128491 | Mammalian PhastCons conservation score<br>https://hgdownload.soe.ucsc.edu/goldenPath/hg19/phastCons46way/placentalMammals/ (hg19)<br>https://hgdownload.soe.ucsc.edu/goldenPath/hg38/phastCons30way/hg38.30way.phastCons/ (hg38) |
| GerpRS | 1064.06/ 1441.64 | GERP++ element score<br>http://mendel.stanford.edu/SidowLab/downloads/gerp/hg19.GERP_elements.tar.gz (hg19)<br>From CADD v1.3 (hg38) |
| GerpRSpv | 1.0/1.0 | GERP++ element p-Value<br>http://mendel.stanford.edu/SidowLab/downloads/gerp/hg19.GERP_elements.tar.gz (hg19)<br>From CADD v1.3 (hg38) |
| rareVar | 0/0 | Number of rare 1KG variants (≤ 5% AF) in a window of ±500 bp<br>http://ftp.1000genomes.ebi.ac.uk/vol1/ftp/data_collections/1000G_2504_high_coverage/working/20200515_EBI_Freebayescalls (hg38) |
| commonVar | 0/0 | Number of common 1KG variants (> 5% AF) in a window of ±500 bp<br>http://ftp.1000genomes.ebi.ac.uk/vol1/ftp/data_collections/1000G_2504_high_coverage/working/20200515_EBI_Freebayescalls (hg38) |
| fracRareCommon | 0.0/0.0 | Ratio rare to common variants<br>http://ftp.1000genomes.ebi.ac.uk/vol1/ftp/data_collections/1000G_2504_high_coverage/working/20200515_EBI_Freebayescalls (hg38) |
| ISCApath | 0/0 | Overlapping ISCA CNVs (date 11/03/2021)<br>https://ftp.ncbi.nlm.nih.gov/pub/dbVar/data/Homo_sapiens/by_study/tsv/nstd46.GRCh37.variant_call.tsv.gz (hg19)<br>https://ftp.ncbi.nlm.nih.gov/pub/dbVar/data/Homo_sapiens/by_study/tsv/nstd75.GRCh37.variant_call.tsv.gz (hg19)<br>https://ftp.ncbi.nlm.nih.gov/pub/dbVar/data/Homo_sapiens/by_study/tsv/nstd102.GRCh37.variant_call.tsv.gz (hg19)<br>https://ftp.ncbi.nlm.nih.gov/pub/dbVar/data/Homo_sapiens/by_study/tsv/nstd46.GRCh38.variant_call.tsv.gz (hg38)<br>https://ftp.ncbi.nlm.nih.gov/pub/dbVar/data/Homo_sapiens/by_study/tsv/nstd75.GRCh38.variant_call.tsv.gz (hg38)<br>https://ftp.ncbi.nlm.nih.gov/pub/dbVar/data/Homo_sapiens/by_study/tsv/nstd102.GRCh38.variant_call.tsv.gz (hg38) |
| dbVARCount | 0/0 | Overlapping dbVAR CNVs (date 10/20/2021)<br>https://ftp.ncbi.nlm.nih.gov/pub/dbVar/archive/Homo_sapiens/by_assembly/GRCh37/gvf/GRCh37.2021_10_20.variant_call.clinical.pathogenic_or_likely_pathogenic.gvf.gz (hg19)<br>https://ftp.ncbi.nlm.nih.gov/pub/dbVar/archive/Homo_sapiens/by_assembly/GRCh38/gvf/GRCh38.2021_10_20.variant_call.clinical.pathogenic_or_likely_pathogenic.gvf.gz (hg38) |

| | | |
|---|---|---|
| DGVCount | 0/0 | Overlapping DGV CNVs (date 02/25/2020)<br>http://dgv.tcag.ca/dgv/docs/GRCh37_hg19_variants_2020-02-25.txt (hg19)<br>http://dgv.tcag.ca/dgv/docs/GRCh38_hg38_variants_2020-02-25.txt (hg38) |

**Supplementary Table 3: ReMM score v0.4 performance** – Area under the precision recall curve (AUPRC) and area under the receiver-operating characteristic curve (AUROC) for ReMM score v0.4 on both genome builds, as well as average values (avg) with standard deviation in parentheses across 100 training runs. AUPRC and AUROC are computed via 10-fold cytoband cross-validation.

| | **GRCh38** | | **GRCh37** | |
|---|---|---|---|---|
| | **Avg of 100 runs** | **ReMM score v0.4** | **Avg of 100 runs** | **ReMM score v0.4** |
| AUPRC | 0.613 (±0.005) | 0.610 | 0.384 (±0.014) | 0.394 |
| AUROC | 0.996 (±0.000) | 0.996 | 0.993 (±0.000) | 0.993 |

**Supplementary Table 4: ReMM score correlation across genome builds** – Pearson and Spearman correlation of ReMM scores between genome builds of three genic regions (DLK1, HBB, PRDM9) and 120,000 random positions (120K). For 120K, only variants with a successful coordinate liftOver from GRCh38 to GRCh37 located on major human chromosomes are used (n=110,751).

| **Name** | **Length (bps)** | **GRCh37 coordinates** | **GRCh38 coordinates** | **Pearson correlation** | **Spearman correlation** |
|---|---|---|---|---|---|
| DLK1 | 123,945 | chr14:101,118,632-101,242,576 | chr14: 100,652,295-100,776,239 | 0.717 | 0.740 |
| HBB | 160,600 | chr11:5,167,199-5,327,798 | chr11:5,145,969-5,306,568 | 0.829 | 0.825 |
| PRDM9 | 106,589 | chr5:23,464,354-23,570,942 | chr5:234,64,245-23,570,833 | 0.685 | 0.692 |
| 120K | 110,751 | - | - | 0.774 | 0.779 |

**Supplementary Table 5: Feature value correlations across genome builds for regions and variants** – Average Pearson and Spearman correlation of the 26 features used for each genome build in three genic regions (DLK1, HBB, PRDM9) and 120,000 random positions (120K). For 120K, only variants with a successful coordinate liftOver from GRCh38 to GRCh37 located on major human chromosomes are used (n=110,751).

| **Name** | **Length (bps)** | **GRCh37 coordinates** | **GRCh38 coordinates** | **Pearson correlation** | **Spearman correlation** |
|---|---|---|---|---|---|
| DLK1 | 123,945 | chr14:101,118,632-101,242,576 | chr14:100,652,295-100,776,239 | 0.680 | 0.686 |
| HBB | 160,600 | chr11:5,167,199-5,327,798 | chr11:5,145,969-5,306,568 | 0.713 | 0.708 |
| PRDM9 | 106,589 | chr5:23,464,354-23,570,942 | chr5:234,64,245-23,570,833 | 0.763 | 0.738 |
| 120K | 110,751 | - | - | 0.572 | 0.650 |

**Supplementary Table 6: ReMM performance dependent on missing values** – Average area under the precision recall curve (AUPRC) and area under the receiver-operating characteristic curve (AUROC) values with standard deviation in parentheses from 100 model training runs using zero as missing values or the default values listed in Supplementary Table 2. AUPRC and AUROC are computed via ten-fold cytoband cross-validation.

| | **GRCh38** | | **GRCh37** | |
|---|---|---|---|---|
| | **Zero** | **Default value** | **Zero** | **Default value** |
| AUPRC | 0.594 (±0.007) | 0.613 (±0.005) | 0.379 (±0.015) | 0.384 (±0.014) |
| AUROC | 0.996 (±0.000) | 0.996 (±0.000) | 0.993 (±0.000) | 0.993 (±0.000) |

**Supplementary Table 7: Feature importance** – Average feature importance (Gini index) over 100 Random Forest partitions of the hyperSMURF models of ReMM v0.4.hg19 and v0.4.hg38. Gini index values were derived with the Ranger package after training on all training data. The standard deviation (std), the minimum (min) and the maximum (max) value across the 100 partitions is shown. The rank indicates the importance rank by average Gini index.

| | | ReMM v0.4.hg38 | | | | | ReMM v0.4.hg19 | | | | |
|---|---|---|---|---|---|---|---|---|---|---|---|
| **Category** | **Feature** | **rank** | **mean** | **std** | **min** | **max** | **rank** | **mean** | **std** | **min** | **max** |
| Conservation | GerpRS | 4 | 0.0258 | 0.0074 | 0.0119 | 0.0444 | 5 | 0.0287 | 0.0082 | 0.0134 | 0.0538 |
| Conservation | mamPhastCons | 7 | 0.0149 | 0.0051 | 0.0013 | 0.0254 | 25 | 0.0000 | 0.0000 | -0.0001 | 0.0002 |
| Conservation | mamPhyloP | 10 | 0.0118 | 0.0048 | 0.0031 | 0.0287 | 10 | 0.0096 | 0.0037 | 0.0022 | 0.0203 |
| Conservation | GerpRSpv | 16 | 0.0036 | 0.0020 | 0.0008 | 0.0134 | 17 | 0.0020 | 0.0014 | -0.0004 | 0.0114 |
| Conservation | verPhyloP | 18 | 0.0033 | 0.0018 | 0.0002 | 0.0103 | 15 | 0.0035 | 0.0018 | 0.0007 | 0.0133 |
| Conservation | priPhyloP | 20 | 0.0020 | 0.0023 | -0.0002 | 0.0141 | 13 | 0.0051 | 0.0029 | 0.0003 | 0.0136 |
| Conservation | verPhastCons | 21 | 0.0018 | 0.0008 | 0.0001 | 0.0048 | 16 | 0.0021 | 0.0010 | 0.0006 | 0.0055 |
| Conservation | priPhastCons | 25 | 0.0008 | 0.0018 | -0.0001 | 0.0177 | 14 | 0.0049 | 0.0031 | 0.0005 | 0.0186 |
| Epigenetic | EncH3K4Me1 | 12 | 0.0106 | 0.0032 | 0.0058 | 0.0186 | 8 | 0.0113 | 0.0031 | 0.0059 | 0.0217 |
| Epigenetic | EncH3K4Me3 | 13 | 0.0105 | 0.0103 | 0.0005 | 0.0551 | 22 | 0.0009 | 0.0012 | -0.0002 | 0.0078 |
| Epigenetic | DnaseClusteredHyp | 14 | 0.0091 | 0.0049 | 0.0004 | 0.0281 | 21 | 0.0012 | 0.0009 | -0.0001 | 0.0062 |
| Epigenetic | EncH3K27Ac | 19 | 0.0030 | 0.0028 | -0.0001 | 0.0121 | 26 | 0.0000 | 0.0000 | -0.0001 | 0.0001 |
| Epigenetic | DnaseClusteredScore | 24 | 0.0014 | 0.0013 | 0.0000 | 0.0087 | 19 | 0.0015 | 0.0008 | -0.0005 | 0.0042 |
| Population | dbVARCount | 1 | 0.1065 | 0.0203 | 0.0672 | 0.1652 | 1 | 0.1062 | 0.0159 | 0.0679 | 0.1385 |
| Population | commonVar | 2 | 0.0657 | 0.0177 | 0.0271 | 0.1044 | 3 | 0.0362 | 0.0140 | 0.0070 | 0.0746 |
| Population | fracRareCommon | 5 | 0.0167 | 0.0063 | 0.0047 | 0.0383 | 6 | 0.0252 | 0.0076 | 0.0116 | 0.0499 |
| Population | rareVar | 9 | 0.0124 | 0.0059 | 0.0023 | 0.0318 | 2 | 0.0366 | 0.0140 | 0.0082 | 0.0875 |
| Population | DGVCount | 15 | 0.0083 | 0.0044 | 0.0008 | 0.0201 | 7 | 0.0115 | 0.0052 | 0.0029 | 0.0293 |
| Population | ISCApath | 17 | 0.0035 | 0.0020 | 0.0002 | 0.0086 | 24 | 0.0007 | 0.0007 | -0.0002 | 0.0039 |
| Regulatory | encRegTfbsClustered | 6 | 0.0164 | 0.0074 | 0.0046 | 0.0390 | 4 | 0.0331 | 0.0120 | 0.0090 | 0.0664 |
| Regulatory | Fantom5Robust | 22 | 0.0015 | 0.0015 | 0.0003 | 0.0101 | 18 | 0.0017 | 0.0009 | 0.0004 | 0.0051 |
| Regulatory | Fantom5Perm | 23 | 0.0014 | 0.0008 | 0.0002 | 0.0053 | 20 | 0.0012 | 0.0014 | -0.0001 | 0.0116 |
| Sequence | CpGperGC | 3 | 0.0339 | 0.0106 | 0.0073 | 0.0583 | 9 | 0.0101 | 0.0054 | 0.0004 | 0.0227 |
| Sequence | CpGperCpG | 8 | 0.0149 | 0.0076 | 0.0038 | 0.0394 | 12 | 0.0066 | 0.0043 | 0.0002 | 0.0206 |
| Sequence | CpGobsExp | 11 | 0.0111 | 0.0046 | 0.0002 | 0.0227 | 11 | 0.0081 | 0.0048 | 0.0004 | 0.0226 |

| | | | | | | | | | | |
|---|---|---|---|---|---|---|---|---|---|---|
| Sequence | GCContent | 26 | 0.0006 | 0.0004 | -0.0001 | 0.0017 | 23 | 0.0009 | 0.0005 | 0.0000 | 0.0024 |

### Supplementary Figures

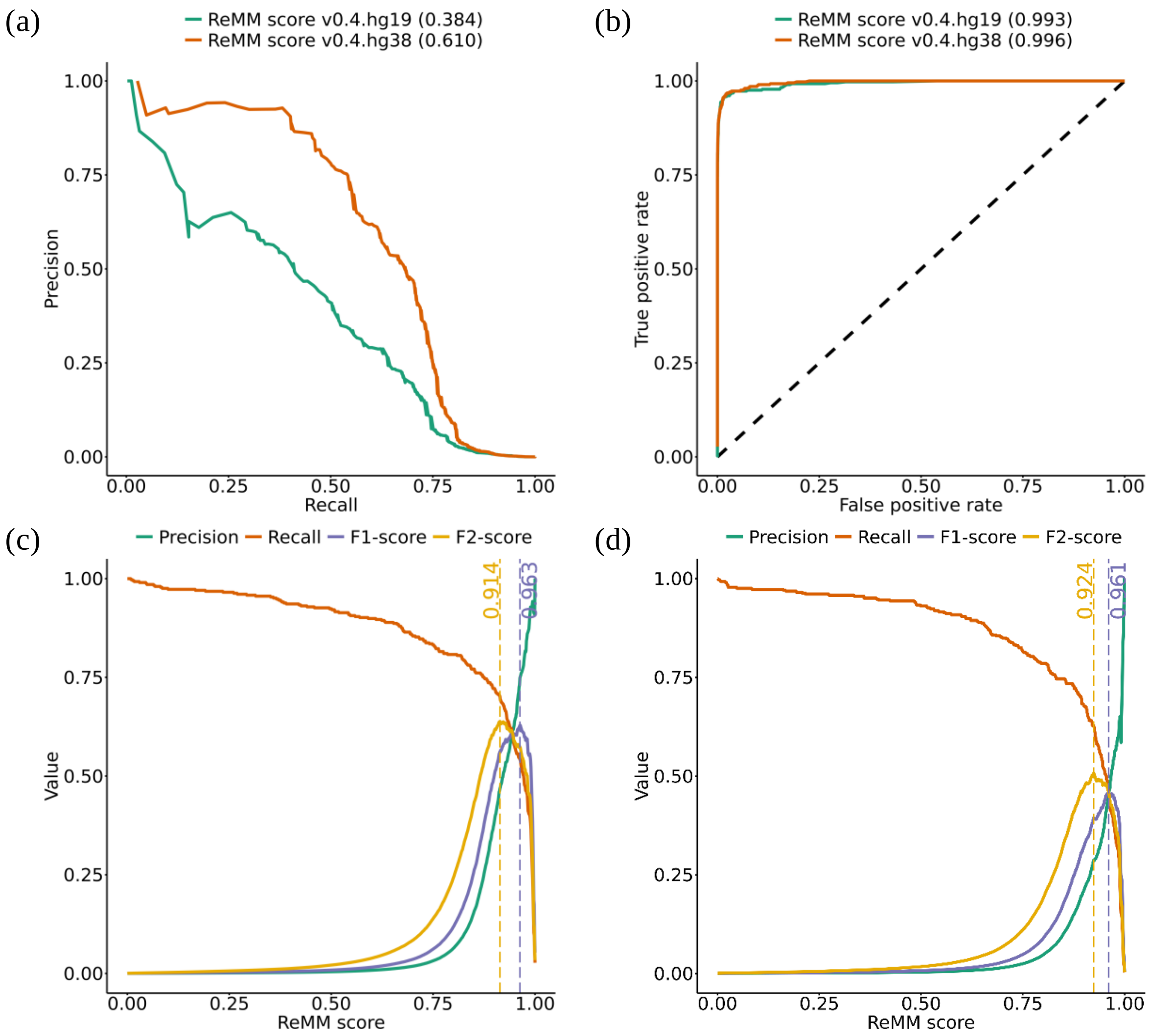


**Supplementary Figure 1: Precision, Recall, ROC, F1-, and F2-score curves** – Performance metric of ReMM v0.4.hg19 and v0.4.hg38 generated via ten-fold cytoband cross validation. Precision-Recall curves (a), receiver operating characteristic curve (ROC) curve (b), and precision, recall F1-score and F2-score (y-axis) over different ReMM score thresholds (x-axis) for v0.4.hg38 (c) and v0.4.hg19 (d). Vertical lines denote the ReMM score with the maximum F1-score (yellow) and the maximum F2-score (purple). Area under the curve is shown in parentheses.

(a) (b)

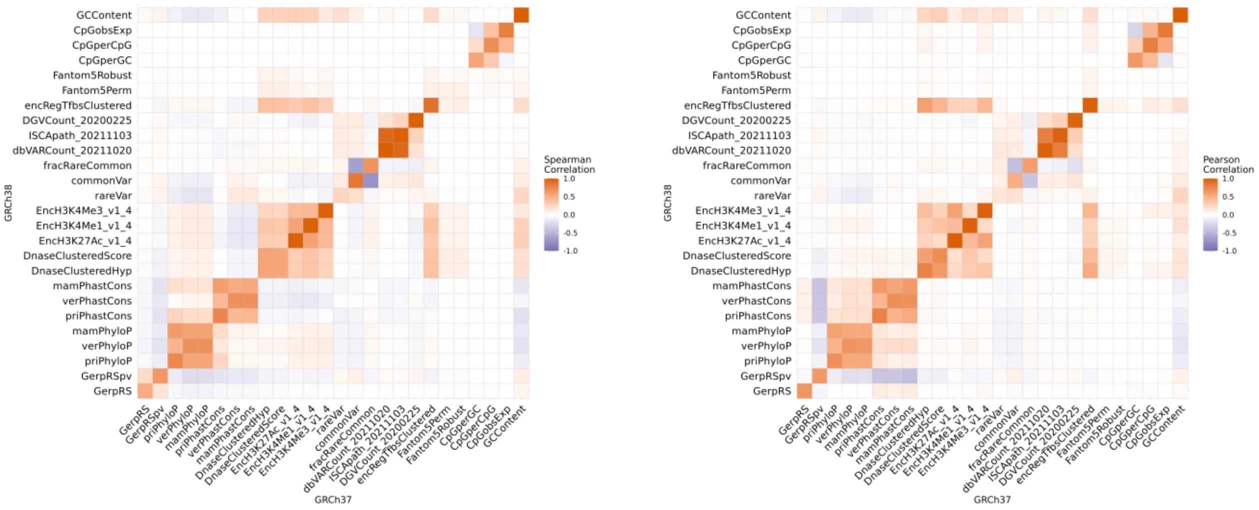


**Supplementary Figure 2: Correlation of feature values**

Feature correlation between features of GRCh37 (x-axis) and GRCh38 (y-axis). The left heat map (a) shows Spearman correlations, and the right (b) shows Pearson correlations. Both plots show unexpectedly low correlations for some features along the diagonal. For example, the histone modification features (ENCODE) are lowly correlated as well as enhancer features (FANTOM).
